# BGX: A Comprehensive Pipeline for Genomic Insight into Bioactivity Prediction, Genomic Surveillance, and Novel Biosynthetic Gene Cluster Assessment

**DOI:** 10.64898/2026.07.20.739398

**Authors:** Ardhendu Chakrabortty, Lovepreet Singh, Babanpreet Kaur, Sunaina Paliyal, Shrikant S Mantri

## Abstract

The increasing availability of genomic and metagenomic data has created significant opportunities to explore microbial diversity, biosynthetic potential and functional traits. However, comprehensive and comparative genome analysis often requires integrating multiple independent tools, making large-scale studies challenging to implement and manage. Here, we present **B**acterial **G**enome e**X**plorer (BGX), an integrated and scalable pipeline that streamlines large-scale genome analysis and exploration of biosynthetic potential. BGX integrates different analytical tools into six major stages: (i) genome retrieval and assembly, (ii) genome quality assessment, (iii) antimicrobial resistance (AMR) gene profiling, (iv) annotating Biosynthetic gene clusters (BGCs) and novelty assessment, (v) bioactivity predictions, and (vi) clustering and networking analysis. In addition, BGX provides a user-friendly interactive interface to facilitate data exploration and interpretation. We demonstrated the versatility and scalability of BGX through large-scale analysis of two independent datasets: 248 genomes from the One Day One Genome (ODOG) initiative and 153 publicly available genomes from NCBI. This analysis enabled the comprehensive characterisation of genome quality, AMR determinants, biosynthetic potential and candidate bioactive metabolites across two datasets. BGX is distributed as a Docker container that simplifies installation, enables reproducible data processing, and supports pipeline execution across different computational environments. The modular and reproducible architecture of the BGX pipeline provides an effective framework for large-scale genome mining, genomic surveillance, and accelerates the discovery and prioritisation of novel secondary metabolites. BGX is now accessible at https://bgx.nabi.res.in

## Introduction

Rapid advances in public datasets and high-throughput sequencing (HTS) technologies have ushered in a new era of biological exploration over the last decade, resulting in a massive and ever-expanding reservoir of genomic data that often exceeds current analytical capacities (Fullam et al., 2023). While the large amount of data provides unprecedented opportunities, its inherent complexity requires highly systematic, directed analytical frameworks to move beyond observational analysis and drive actionable functional interpretations (Malviya et al., 2022). Intricate analysis of such datasets helps to glean functional insights and uncover the metabolic potential of microbial communities (Mandal et al., 2022). Therefore, the interpretation of previously uncharacterized secondary metabolites by such analysis plays a crucial role.

The directed functional annotation and comparative analysis of genomic datasets form a critical bridge between complex biological interpretations and raw sequencing data (Gavriilidou et al., 2022). Over a decade, the synergistic evolution of computational tools and databases has significantly improved their performance (Ziemert et al., 2016). A highly prized outcome of such genome mining is the interpretation of novel BGCs (Blin et al., 2023; Cimermancic et al., 2014). In brief, BGCs are co-localized genes structurally in microbial genomes that coordinate to encode the biosynthetic pathways for producing secondary metabolites (SMs) and their chemical variants (Cimermancic et al., 2014; Medema et al., 2015). These specialized co-localized genomic regions function as a layout for SMs biosynthesis, empowering exploration of uncharacterized natural products from a vast repertoire generated by microbial communities inhabiting diverse and competitive niches (Crits-Christoph et al., 2018). Moreover, the robust BGC prediction and profiling facilitate identification and prioritization of high-potential microbial taxa and specific functional pathways as targets for metabolite isolation and downstream experimental validation (Katz & Baltz, 2016). Ultimately, this targeted approach streamlines the discovery of potential natural products produced by these microbial communities.

There are dual functions supported by the increasing abundance of publicly available genomic and metagenomic datasets. Majorly, these global datasets are important for systematic genome mining for the interpretation of a vast array of natural products encoded within the complex microbial community (Struelens et al., 2024; Zhang et al., 2024). Also, the in-depth analysis of datasets enables the researcher to explore the emerging resistance patterns and mechanisms driving antimicrobial resistance (AMR). The AMR pandemic has emerged as a critical global challenge that goes beyond human health (Graham et al., 2016; Liu et al., 2016). Natural reservoirs of antibiotic biosynthesis are the competitive ecological niches, and resistance mechanisms co-evolve in dynamic evolution (Crits-Christoph et al., 2018). Not only does interpretation of natural products, but also surveillance of AMR spread across taxa, and precise functional analysis (Hendriksen et al., 2019).

To analyse large and complex genomic datasets, a wide range of computational tools and integrated workflows have been developed for BGC discovery and interpretation. Some approaches integrate taxonomic information with genome mining to improve the detection of experimentally characterised BGCs and infer secondary metabolite production potential in metagenomic datasets. (Gupta et al., 2022). Others focus on quantifying the abundance and transcriptional activity of BGCs across microbiomes using high-throughput sequencing data, thereby enabling functional characterisation of microbial communities. (Pascal Andreu et al., 2021) (Pascal Andreu et al., 2021). Recent workflows have expanded genome mining to pangenome-scale analyses, allowing systematic exploration of BGC diversity across thousands of genomes, while long-read sequencing–oriented approaches improve assembly continuity and facilitate the recovery of complete gene clusters from complex samples. Comparative genomics frameworks further enable the identification of homologous BGC families across related taxa, providing insights into their evolutionary conservation, diversification, horizontal gene transfer, and the discovery of novel variants from metagenomic data (Salamzade et al., 2023). Together, these tools and workflows aim to streamline genome mining processes, enabling scalable, reproducible, and high-throughput analysis of BGCs across diverse datasets (Agrawal et al., 2026; Nuhamunada et al., 2024; Salamzade et al., 2023).

In this study, we present BGX, a comprehensive and reproducible workflow for large-scale bacterial genome analysis. The primary aim of BGX is to provide an integrated framework that enables the systematic characterization of bacterial genomes by combining multiple analytical modules within a single, automated pipeline. In addition to BGCs discovery and comparative analysis, BGX supports the identification of antimicrobial resistance (AMR), the detection of C-type lectin proteins, and the prediction of bioactivity. BGX integrates diverse analytical steps into a unified workflow, thereby minimizing fragmented tool usage, reducing manual intervention, and simplifying the execution of complex analyses. The pipeline also addresses common challenges associated with bioinformatics workflows, such as software version incompatibilities, dependency conflicts, and database mismatches, enabling scalable and reproducible analyses across diverse computational environments.

By integrating these complementary modules, BGX provides deeper insights into the biosynthetic potential, resistance profiles, and lectin repertoires of bacterial genomes, thereby supporting comprehensive genomic investigations. To demonstrate the applicability and robustness of the BGX pipeline, it was applied to representative genomic datasets encompassing diverse environmental and genomic contexts. These use cases highlight the pipeline’s ability to process heterogeneous inputs and generate comprehensive insights into BGCs, AMR genes, and C-type lectins in bacterial genomes.

## Methods

### Data retrieval and genome assembly

To demonstrate the applicability and versatility of BGX, genomes obtained from three different sources were analysed: (i) genome assemblies downloaded using the NCBI Datasets tool, (ii) genome sequences retrieved from the NCBI Nuccore database using efetch, and (iii) raw sequencing reads, which were assembled de novo using SPAdes (v3.15.5) to generate contigs (Prjibelski et al., 2020). For quality check of the assembled genomes BUSCO (v6.0.0) was used to identify complete, single copy, duplicate, fragmented, and missing copies. As a lineage database, we used “bacteria_odb10”. The analysis was performed with genome mode on.

### BGCs Identification

BGCs were identified using a combination of tools, including antiSMASH (v8), DeepBGC, and GECCO, to ensure comprehensive and sensitive detection of BGCs (Blin et al., 2025; Carroll et al., 2021; Hannigan et al., 2019). antiSMASH analysis was performed with a minimum cluster length threshold of 1000 bp. antiSMASH identifies BGCs based on signature genes, domain architecture, and rule-based detection of cluster classes such as non-ribosomal peptide synthetases (NRPSs), polyketide synthases (PKSs), ribosomally synthesized and post-translationally modified peptides (RiPPs), and terpenes. DeepBGC, which uses deep learning, and GECCO, which leverages machine learning for gene cluster prediction, were used to complement antiSMASH predictions and improve the detection of novel and atypical BGCs (Blin et al., 2025; Carroll et al., 2021; Hannigan et al., 2019).

### BGC clustering

To assess the sequence and architectural diversity of the predicted BGCs, Biosynthetic Gene Similarity Clustering and Prospecting Engine v2 (BiG-SCAPE) (Draisma et al., 2026) was employed to group BGCs into gene cluster families (GCFs) based on domain composition and sequence similarity. The BiG-SCAPE run was performed with a default cutoff of 0.3, using BGCs from the Minimum Information about a Biosynthetic Gene cluster (MIBiG) database v4 as the reference (Zdouc et al., 2025). The resulting similarity networks were analyzed to identify relationships among clusters and evaluate their relatedness to previously characterized biosynthetic systems. This analysis facilitated exploration of BGC diversity, identification of potentially novel cluster families, and prioritization of candidate BGCs for downstream investigation.

### Bioactivity predictions

To gain insights into the potential bioactivities associated with the annotated BGCs, a neural network-based bioactivity prediction tool, NPBdetect, was utilized. This tool can detect multiple bioactivities in cluster sequences, including antibacterial, antifungal, cytotoxic/antitumor, siderophore, antiviral, antiprotozoal, inhibitory, and surfactant activities (Goyat et al., 2025). BGCs obtained from antiSMASH (v8) were used as input and predicted to be bioactive positive with a probability score ≥ 0.5.

Further to increase the reliance on the predicted bioactivities, we employed another prediction tool, natural product function (NPF), which is currently not part of our pipeline. NPF is a machine learning model that can detect antibacterial, antifungal, and cytotoxic/antitumor bioactivities (Walker & Clardy, 2021). An overlap analysis was performed between bioactive BGCs predicted by NPF and NPBdetect to identify clusters associated with antibacterial, antifungal, and cytotoxic/antitumor activities. This approach enabled the identification of high-confidence bioactive BGCs supported by concordant predictions from both tools, thereby improving overall prediction reliability.

PROkaryotic DYnamic programming Gene-finding ALgorithm (Prodigal) (v 2.6.3) (Hyatt et al., 2010) or Prokka (v 1.12) (Seemann, 2014) were used to convert genomes to proteins and used pHMMS to search for C-type lectins using the program hmmsearch part of HMMER (Finn et al., 2011). We also did NCBI-BLASTp analysis of the found proteins. C-type lectin prediction ML tool is also available (Singh et al., 2024).

### Siderophoric potential

NPBdetect predicted siderophore-positive BGCs and their corresponding bacterial genomes were further screened for siderophore-associated transporter domains, including FecCD, periplasmic binding proteins (PBP), and TonB-dependent receptors (TonB), using an additional tool HMMER v3.3.2, which is currently not part of this pipeline (Potter et al., 2018), following previously reported domain associations (Crits-Christoph et al., 2021; Reitz & Medema, 2022). These predicted BGCs were subsequently evaluated against antiSMASH siderophore detection rules to identify biosynthetic features linked to known siderophore structural classes.

Further, the sequence similarity network was constructed for the predicted siderophore BGCs with 78 reference NRP-metallophore BGCs (Reitz et al., 2026) to identify the core siderophore moieties associated with the predicted siderophore BGCs.

### BLASTn analysis against SMC database

The predicted BGC sequences were subjected to BLASTn analysis against the secondary metabolism collaboratory (SMC) nucleotide database to determine whether they corresponded to previously characterized BGCs or represented putatively novel clusters (Camacho et al., 2009; Udwary et al., 2025). For each query BGC, only the best hit, defined as the match with the highest query coverage (qcov), was retained for downstream analysis. Based on qcov values, BGCs were classified into three categories: KNOWN_LIKE (qcov ≥ 80%), RELATED (40% ≤ qcov < 80%), and NOVEL (qcov < 40%), according to the classification scheme adopted in this study.

### Antibiotic resistance profiling

The resistance gene identifier (RGI) (v6.0.5) (Alcock et al., 2023) was used to predict AMR using the comprehensive antibiotic resistance database for both use cases. RGI from bioconda was installed and run locally within the pipeline, and the predicted resistance was profiled using the rgi in-built heatmap feature. A prevalence parameter of 10% was used to determine the prevalent AMR genes. AMR gene overlaps across sources were plotted with the help of ggvenn R package (Yan, 2025).

### Pipeline Implementation

BGX pipeline is distributed as an open-source GitHub repository together with a containerized Docker image containing all required software packages, databases, and workflow dependencies. Users can execute the complete pipeline directly from the Docker container without manually installing individual tools or resolving software compatibility issues. This containerized implementation ensures portability, simplifies deployment across different computing environments, and improves the reproducibility of genomic analyses.

## Results

### Pipeline architecture and performance

BGX combines multiple state-of-the-art tools and analytical modules for BGCs prediction, novelty assessment against reference databases, comparative clustering of related gene clusters, bioactivity inference, AMR gene identification, and C-type lectin detection (Figure 1). To demonstrate the robustness and applicability of the BGX pipeline, we present its performance across two independent use cases: (i) an “One Day One Genome” (ODOG) dataset of 248 Indian genomes from the ODOG initiative, launched by the Biotechnology Research and Innovation Council (BRIC), India; (ii) an Indian deposited NCBI dataset (IndeNCBI) of 153 publicly available genomes. These use cases are presented independently to highlight the versatility of the pipeline in handling datasets.

**Figure 1.**
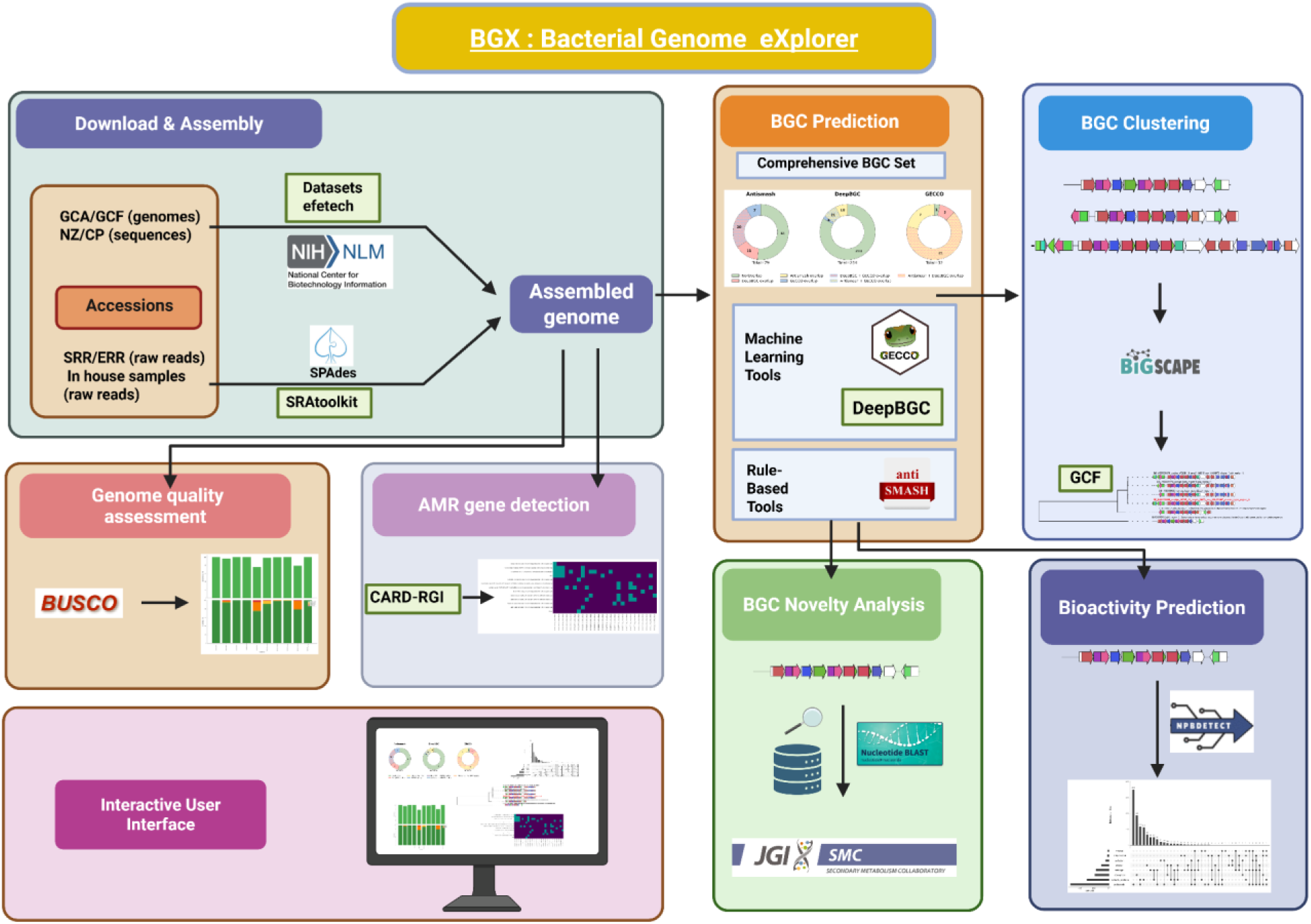
A schematic representation of the BGX workflow. Input genomic data (GCA/GCF assemblies or SRR/ERR reads) are processed through genome assembly (SPAdes) followed by BGC prediction using antiSMASH, DeepBGC, and GECCO. Downstream analyses include BGC novelty assessment (e.g., comparison to reference databases), clustering (BiG-SCAPE), and bioactivity prediction (e.g., NPBdetect). Genome quality assessment (BUSCO) and antimicrobial resistance gene detection (RGI) are also incorporated, with results visualized through an interactive interface.

For a representative dataset of 10 genomes, BGX completed execution in 8 h 8 min (wall time), utilizing ∼23 h total CPU time (∼3 cores in parallel) with a peak memory usage of ∼57 GB and no observed disk swapping.

### Use case 1: Application of BGX pipeline on the ODOG initiative data analysis

We included 248 genomes from the ODOG initiative of BRIC in India. The ODOG dataset is largely dominated by human-associated samples (∼34.3%), indicating a strong representation of host-associated environments. Among environmental sources, wastewater (∼13.3%) and soil (∼10.9%) are the major contributors, while animal (∼8.1%) and plant-associated (∼7.3%) samples also form a significant portion.

Moderate representation is seen in unknown (∼6%), fermented food (∼5.2%), and food (∼4.4%) categories, while aquatic environments such as sewage, lake, spring, and well water (∼1–2%) contribute minimally (Supplementary Table 1A). Overall, the dataset reflects a diverse habitat distribution, with human- and wastewater-associated samples being the most prominent categories.

### Application of BGX to assess assembly quality

Assembly quality was assessed using BUSCO (v6.0.0) with the bacteria_odb10 dataset in genome mode, showing an average completeness of 95.77%. This indicated a high level of genome completeness among 248 ODOG genomes. On average, assemblies were 90.95% single-copy, 7.41% duplicated BUSCOs, 1.8% fragmented BUSCOs, and 2.6% missing BUSCOs. The assessment categories from BUSCO analysis were presented using a dual-panel plot (Supplemetary Figure 1A). We observed that the majority of the conserved orthologs were present as a single copy. Collectively, the genome assemblies exhibited high quality, rendering them suitable for subsequent downstream analyses.

### Multi-tool BGCs detection using BGX

In the ODOG dataset, antiSMASH predicted 1,944 BGCs. For the overlap analysis, only BGCs larger than 5 kb were retained, as shorter clusters are unlikely to represent meaningful biosynthetic regions and were therefore excluded. DeepBGC predicts the highest number of BGCs (total = 3418), followed by antiSMASH (1797) and GECCO (730). A large proportion of non-overlapping predictions (e.g., 2732 in DeepBGC and 1073 in antiSMASH) highlights the sensitivity of DeepBGC and tool-specific detections (Supplementary Table 1B). In contrast, substantial multi-tool overlaps (∼400 BGCs shared across all three tools) indicate a core set of high-confidence BGCs. This pattern (Figure 3A) reflects greater novelty and diversity in the ODOG dataset, emphasizing the multi-tool integration approach of BGX to capture diverse biosynthetic potential.

The antiSMASH annotated BGCs were further processed for clustering using BiG-SCAPE v2. At 0.3 cutoff, under “mix” category, all the 1,944 BGCs were clustered into 1297 distinct GCFs. 1198 out of 1297 GCFs were considered as novel due to lack of MIBiG reference BGCs whereas the remaining 99 GCFs clustering with known MIBiG reference BGCs were considered as known GCFs (Supplementary Table 1C).

### Biosynthetic space exploration and novelty assessment with BGX

Within BGX, the novelty of BGCs was assessed using BLASTn-based sequence similarity searches against the SMC database. In the ODOG dataset, out of a total of 1944 BGCs, 1372 were classified as KNOWN_LIKE, indicating a substantial proportion of previously characterized clusters (Figure 2). Within this category, RiPPs (366) and other (287) classes are most abundant, followed by NRPS (264), PKS (196), and terpenes (168), along with multiple hybrid combinations. This reflects a broad representation of established biosynthetic pathways (Supplementary Table 1D). The abundance of RiPP BGCs in the ODOG dataset is consistent with previous studies showing that RiPPs are widespread in human-associated microbiomes and contribute to microbial competition and host interactions (Donia et al., 2014). The RELATED category (211 BGCs) shows moderate similarity to known clusters and is distributed across diverse classes, including terpenes (73), other (75), and RiPP (25), with smaller contributions from NRPS and PKS hybrids (Figure 2). This suggests the presence of diverse biosynthetic systems with potential functional divergence. A significant proportion of clusters fall into the NOVEL category (361 BGCs), indicating extensive unexplored biosynthetic space. Notably, this category is strongly dominated by terpenes (241), followed by RiPPs (73) and other classes (30), with only minor representation from the NRPS and PKS groups. The enrichment of putatively novel terpenes and RiPPs suggests that these classes may represent promising sources of previously uncharacterized natural products.

**Figure 2.**
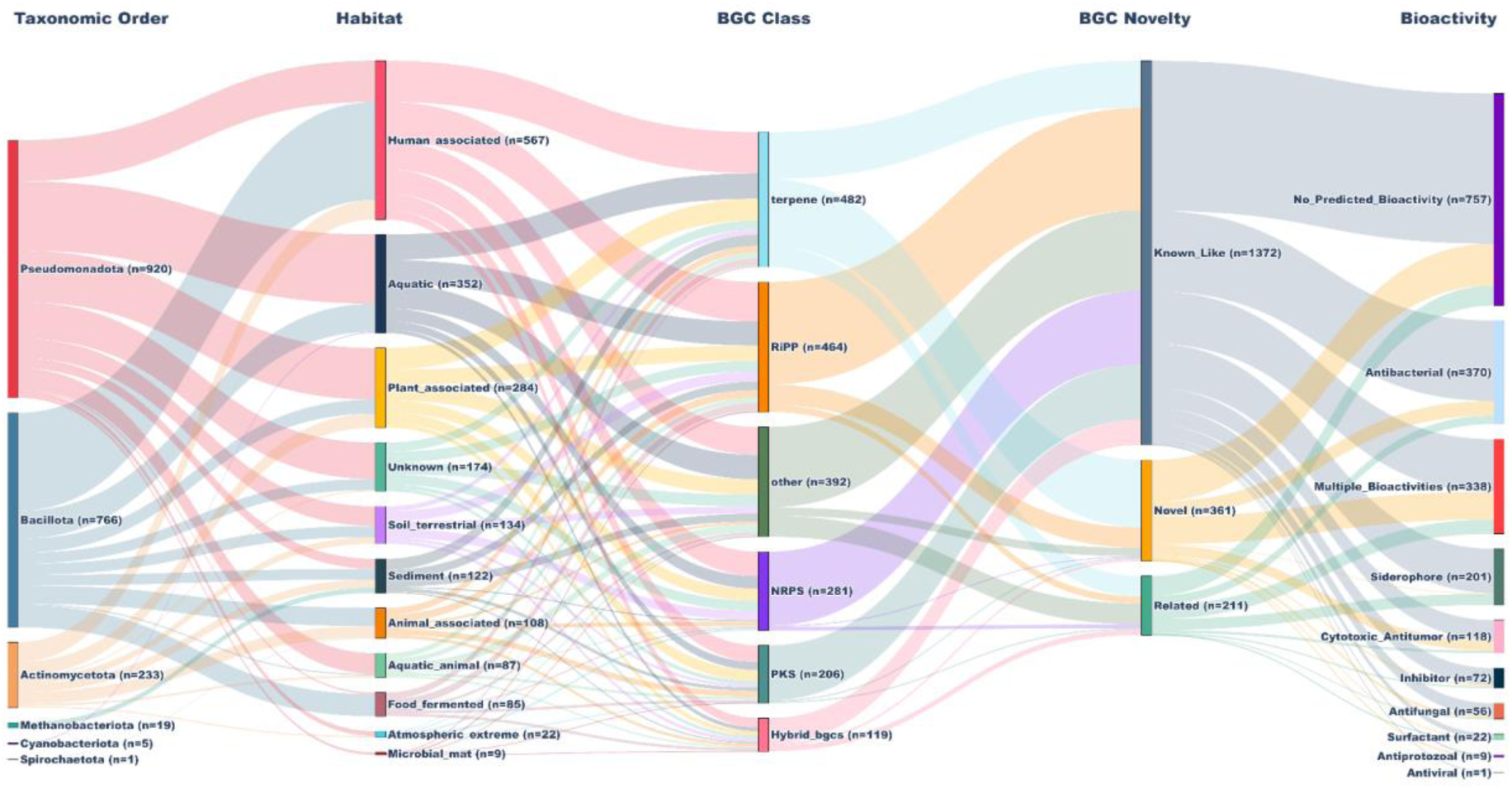
The Sankey diagram illustrates the relationship between environmental habitats, predicted BGC classes, and their novelty status (known-like, related, or novel). It shows how BGCs originating from different habitats are distributed across various biosynthetic classes and further categorized based on their similarity to previously characterized clusters.

### BGX-enabled bioactivity predictions reveal a rich repertoire of antibacterial BGCs

To assess the bioactivity potential of annotated BGCs, BGX uses the neural network-based bioactivity prediction tool NPBdetect (Goyat et al., 2025). Among the 1,944 BGCs, 594 were predicted to encode antibacterial activity, 344 cytotoxic/antitumor activity, 218 siderophore activity, 206 antifungal activity, and 120 inhibitory activity (Supplementary Table 2). To examine the overlap among predicted bioactivities, an UpSet plot (ConFigure 2way et al., 2017) was generated from the prediction output. This analysis showed that of the 594 antibacterial BGCs, 370 were predicted exclusively as antibacterial, while 201 of 218 BGCs were predicted exclusively as siderophores (Figure 3B).

**Figure 3:**
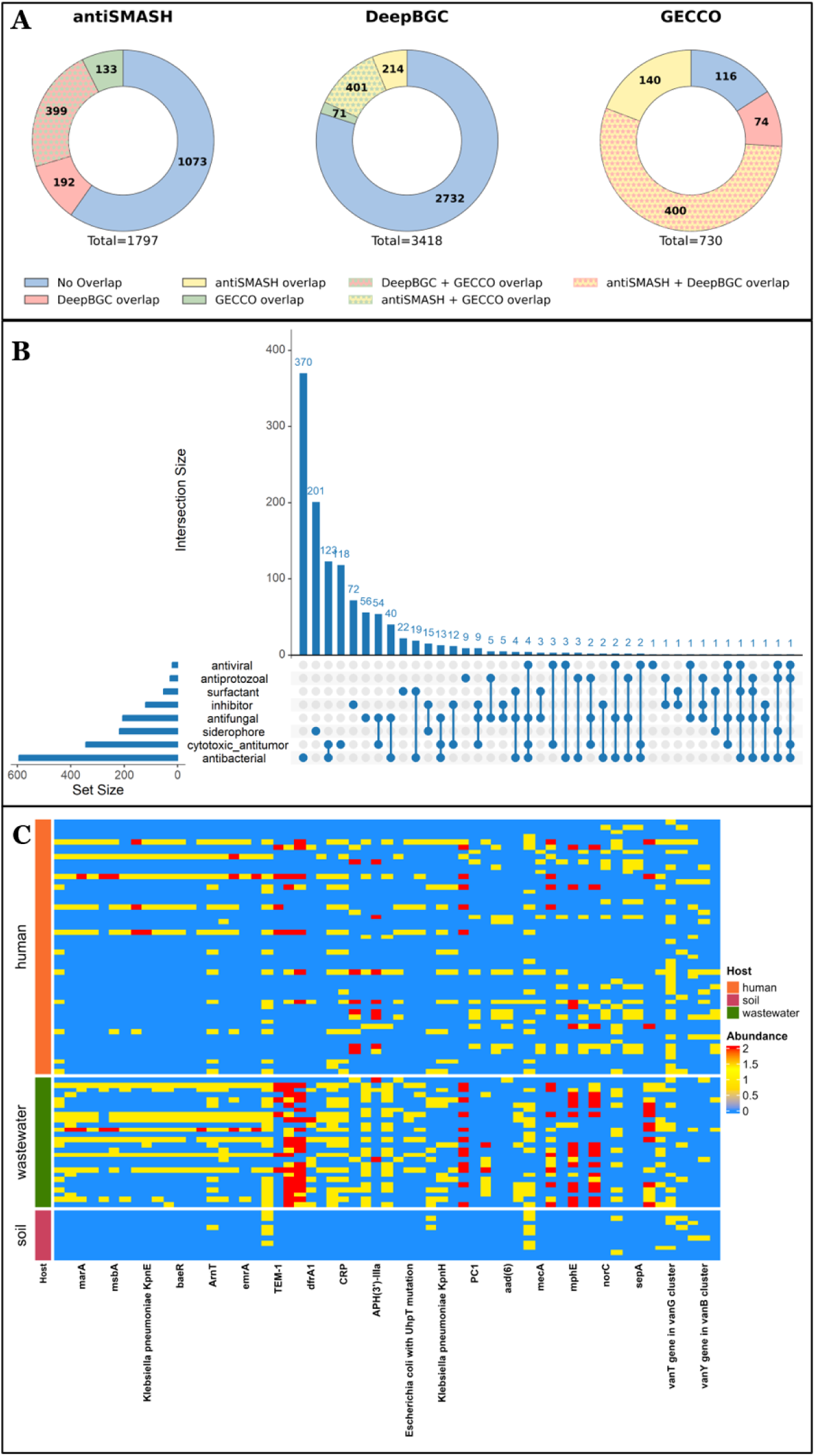
Multipanel figure for ODOG dataset (A) BGC prediction overlap across tools in the ODOG dataset (B) An upset plot showing the overlap of predicted bioactivities of BGC and (C) A heatmap presenting the top 10% prevalent AMR genes distribution across habitat. Donut plots showing the overlap of BGCs predicted by antiSMASH, DeepBGC, and GECCO in the ODOG dataset. Each segment represents either unique predictions (no overlap) or shared predictions between tools (pairwise or multi-tool overlaps). Total BGC counts per tool are indicated below each plot. The vertical bars represent the shared bioactivities of BGCs annotated from the ODOG dataset.

To further improve prediction reliability, another machine learning-based model, NPF, was used to predict antibacterial, antifungal, and cytotoxic/antitumor bioactivities (Walker & Clardy, 2021). An overlap analysis of antibacterial, antifungal, and cytotoxic/antitumor predictions from the NPBdetect and NPF tool was performed. This analysis revealed that 417 BGCs were consistently predicted to be antibacterial, 88 to be cytotoxic/antitumor, and 55 to be antifungal by both prediction tools. Thereby, our analysis pipeline provides greater confidence in the robustness of these bioactivity predictions (Supplementary Figure 1B).

### Characterization of predicted siderophore BGCs to provide additional insights into the existing pipeline

Bioactivity predictions by NPBdetect identified 218 BGCs with predicted probabilities ≥0.5 for siderophore activity. As an addition to this pipeline, we employed HMMER for sequence homology and domain detection. Siderophore domain detection was conducted as an independent analysis, as this feature is not currently integrated into the BGX pipeline. Among these, 144 predicted BGCs were found to encode at least one of the key siderophore-associated transporter domains, namely TonB, FecCD, and PBP. Examination of the domain distribution revealed both distinct and overlapping patterns among these transporter-associated components. The most prevalent architecture comprised the simultaneous presence of all three domains, observed in 52 BGCs, indicating that 36.1% of predicted siderophore BGCs encoded a relatively complete transport and uptake machinery (Supplementary Figure 1C). In addition, 39 BGCs (27.1%) contained the FecCD–PBP combination, while 7 BGCs (4.9%) encoded PBP–TonB and 3 BGCs (2.1%) encoded FecCD–TonB.

Analysis of antiSMASH siderophore-associated detection rules revealed a predominance of NRPS-independent siderophore (NI-siderophore) pathways, accounting for 58.0% of the predicted siderophore BGCs. This was followed by NRP-metallophore-associated clusters, which accounted for 40.1% of the predictions, indicating a substantial contribution from NRPS-dependent biosynthetic mechanisms (Supplementary Figure 1D). In contrast, other siderophore-associated rule types, including opine-like metallophore pathways, were underrepresented (∼1.3%), highlighting their relative rarity. Overall, the distribution suggests that NI-siderophore biosynthetic strategies are the dominant mode within the ODOG dataset, with additional diversity contributed by NRPS-based metallophore systems.

Further, the clustering of predicted siderophore BGCs with reference NRP-metallophore BGCs (Reitz et al., 2026) revealed the clustering of 22 predicted BGCs with reference metallophore BGCs, highlighting the presence of core siderophoric moieties such as hydroxamate, salicylate, OHAsp and graminine, OHAsp and hydroxamate (supplementary Table 3).

### Profiling and surveillance of AMR gene dissemination using BGX pipeline

As major sources of dissemination and the largest reservoirs of AMR genes, genomes isolated from soil, wastewater, and human samples were analysed using CARD-RGI to detect resistance genes. A total of 546 resistance genes were detected, and the number of hits per gene was recorded. We observed notable concordance in the resistance profiles of wastewater and human samples (Supplementary Figure 1E). An overlap of 28.3% (n=126) resistance genes was observed. However, 13 resistance genes, consisting of 2.9%, were present across all three sources. Notably, samples collected from human sources had the highest percentage of unique genes (39.5%, n=176). Likewise, wastewater had the second-highest number of unique resistance genes, 24.9% (n=111), followed by soil, 2.7% (n=12) (Supplementary Figure 1F). However, soil-to-human transmission and soil-to-wastewater transmission remained very low, with an overlap of 0.9% each. We have listed the top ten most prevalent AMR genes across habitats: *rsmA*, *vanT* gene in *vanG* cluster, *adeF*, *sul1*, *ArnT*, Escherichia coli EF-Tu mutants conferring resistance to Pulvomycin, *CRP*, Haemophilus influenzae PBP3 conferring resistance to β-lactam antibiotics, *qacEdelta1*, and *APH(3”)-lb*. The spread of several alarming AMR genes, including *TEM-1*, *APH(3”)-lb, sul1*, *mphA*, and *mphE*, was highlighted by a heatmap representing the distribution of prevalent genes across different habitats (Figure 3C). We observed that 28.3% of resistance genes were shared between wastewater and human samples. As reported before, soil and wastewater are the biggest reservoirs of AMR genes and a major source of dissemination (Das et al., 2025; Islam et al., 2025; Read et al., 2024; Sassi et al., 2025). It is also noticed that, 39.5% resistance genes were unique to human samples in ODOG dataset. This warrants further investgation in to contribution from other habitats, over-the-counter availability of drugs, and improper utilisation of antibiotics. For a country like India, AMR genes were also reported to be circulated within the hospital-ICU environment (Chakrabortty et al., 2024). In our analysis, it came out that *rsmA* is the most prevalent resistance gene, which is a post-transcriptional regulatory protein, controlling several key functions related to virulence, including biofilm formation (Mulcahy et al., 2008). The presence of *ArnT*, the second most prevalent AMR gene, was also alarming, as it offered resistance to the polymyxin class of antibiotics (Sasal et al., 2025). Prevalence of *sul1* and *qacEdelta1* resistance genes was previously reported as prevalent ARG in a surveillance study conducted in Japan (Sekizuka et al., 2022). This highlights the BGX pipeline’s role in tracking resistance gene transmission from the environment like wastewater to humans.

### Use case 2: Implementation of the BGX pipeline for analysis of the IndeNCBI dataset

For the second use case, we included 153 genomes representing Indian isolates deposited in the last 10 years. The IndeNCBI dataset is predominantly composed of food-associated samples (∼33.1%), making it the largest source category. This is followed by animal-associated samples (∼15.6%) and human-associated samples (∼13.6%), indicating a strong representation of host-associated microbiomes. Environmental contributions include soil (∼12.3%), while a notable fraction is attributed to sources unidentified (∼13.0%). Minor representation is seen from plant-associated samples (∼2.6%), natural environments (∼1.9%), and lab stock (∼1.3%) (Supplementary Table 4A). Overall, the dataset shows a more balanced but less diverse distribution, with a strong emphasis on food and host-associated environments compared to environmental niches.

### Application of BGX to assess assembly quality

Assembly quality was assessed using BUSCO (v6.0.0) with the bacteria_odb10 dataset in genome mode, showing an average completeness of 98.50%. This indicated a very high level of genome completeness among 153 IndeNCBI genomes. On average, assemblies were 99.51% single copy, 0.48% duplicated BUSCOs, 0.78% fragmented BUSCOs, and 0.72% of missing BUSCOs. The BUSCO-based genome completeness and assessment categories were visualized using a dual-panel plot (Supplementary Figure 2A). Majority of the conserved orthologs were present as a single copy underscoring the high quality of the genome assemblies and suitability for downstream processing.

### Multi-tool BGCs detection using BGX

In the IndeNCBI dataset, antiSMASH predicted 1,215 BGCs. DeepBGC again identifies the largest number of BGCs (total = 2781), followed by antiSMASH (1210) and GECCO (685), for BGCs >5000 bp. Compared to the index case, the proportion of overlapping predictions is higher (e.g., 408 shared across all three tools along with strong pairwise overlaps), while the number of unique predictions is relatively lower (e.g., 2104 in DeepBGC, 568 in antiSMASH) (Supplementary Table 4B). This trend (Figure 5A) suggests more conserved and well-characterized biosynthetic regions. The higher agreement between tools indicates increased confidence in detected clusters across datasets.

Further clustering of the antiSMASH-annotated 1,215 BGCs was performed using BiG-SCAPE v2 to group them into GCFs based on sequence similarity. At cutoff 0.3, under the “mix” category, 1,215 BGCs from the IndeNCBI dataset were clustered into 498 distinct GCFs. Among these 427 GCFs did not cluster with any MIBiG reference BGCs, whereas the remaining 71 GCFs clustered with MIBiG reference BGCs, highlighting their potential to encode similar known compounds (Supplementary Table 4C).

### Novelty Assessment of BGCs with BGX

The novelty of BGCs identified in the NCBI dataset was assessed using BLASTn-based sequence similarity searches against the SMC reference database. Out of a total of 1215 BGCs, the majority (981 BGCs) were classified as KNOWN_LIKE, indicating strong similarity to previously characterized clusters (Figure 4). Within this category, RiPPs (329) and NRPS (238) dominate, followed by other (153), terpenes (77), and PKS (74), along with several hybrid classes (e.g., NRPS–PKS, PKS–terpene) (Supplementary Table 4D). This highlights the prevalence of well-characterized biosynthetic pathways, particularly peptide-based and ribosomally synthesized clusters. NRPS BGCs are widely distributed in environmentally diverse bacteria, particularly in soil and marine ecosystems, where they encode structurally diverse metabolites involved in microbial competition, nutrient acquisition, and ecological adaptation (Al-Siyabi et al., 2025). A smaller fraction (164 BGCs) was categorized as RELATED, representing moderately similar clusters. This group is strongly enriched in terpene clusters (97), along with contributions from other (33) and RiPP (19) classes, and only minimal representation from NRPS/PKS categories. The NOVEL category comprises 70 BGCs, with a clear dominance of terpenes (58), and very few RiPP (6), other (5), and PKS (1) clusters. This indicates that most of the novel biosynthetic potential in this dataset is concentrated within terpene pathways. Overall, the novelty distribution shows that the NCBI dataset is largely composed of conserved and well-characterized BGCs, with limited but distinct novelty primarily driven by terpene clusters and relatively low representation of novel NRPS and PKS systems.

**Figure 4.**
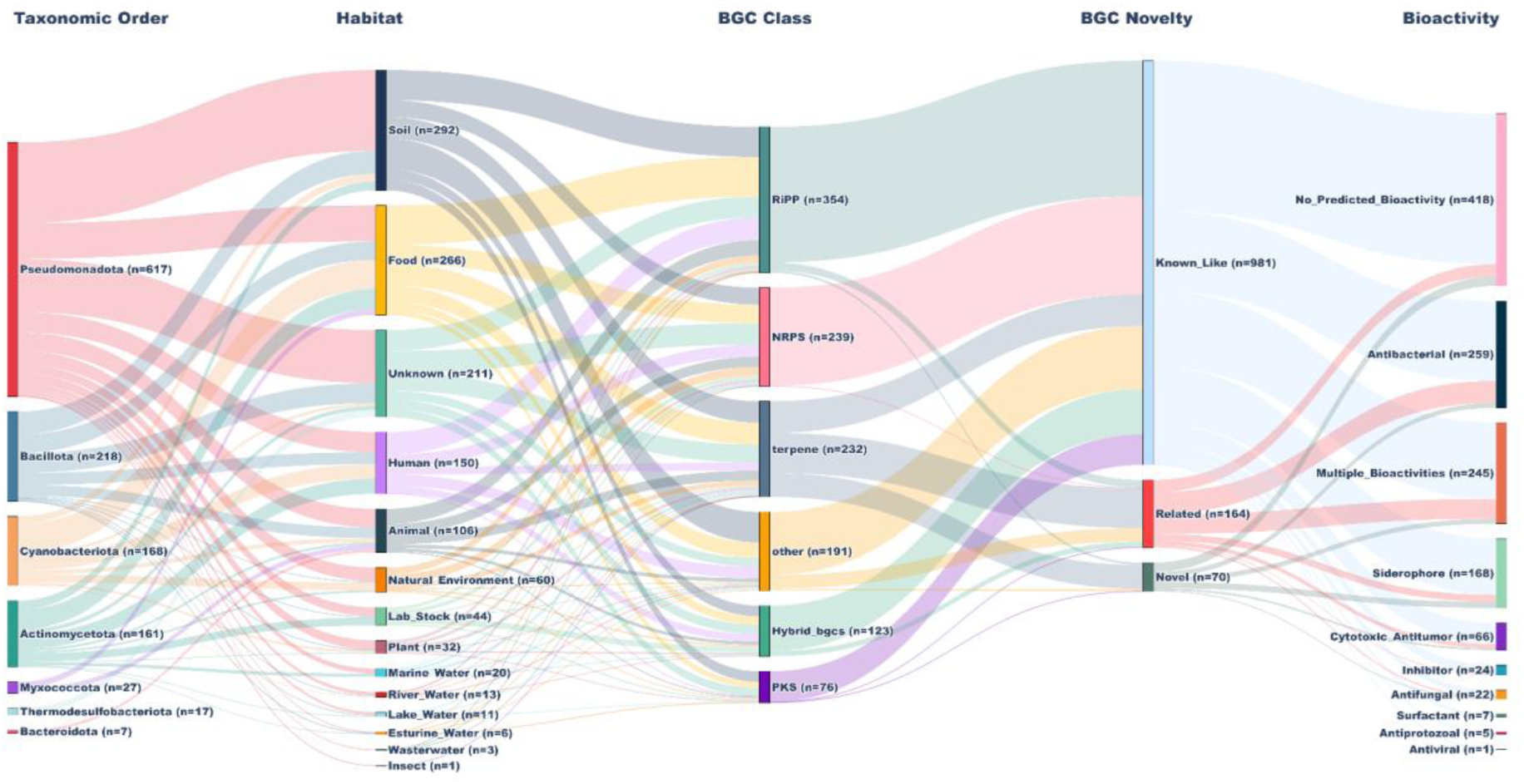
The Sankey diagram illustrates the relationship between environmental habitats, predicted BGC classes, and their novelty status (known-like, related, or novel). It shows how BGCs originating from different habitats are distributed across various biosynthetic classes and further categorized based on their similarity to previously characterized clusters.

### BGX-enabled bioactivity predictions reveal a rich repertoire of antibacterial BGCs

Bioactivity predictions using NPBdetect identified 425 BGCs with predicted probabilities of antibacterial, 248 cytotoxic/antitumor, 180 siderophore, and 117 antifungal bioactivity (Supplementary Table 5). The UpSet plot showcasing the overlap of predicted bioactivities revealed that, of the 425 antibacterial predicted BGCs, 259 were predicted exclusively as antibacterial, while 168 of 180 BGCs were predicted exclusively as siderophore (Figure 5B).

**Figure 5:**
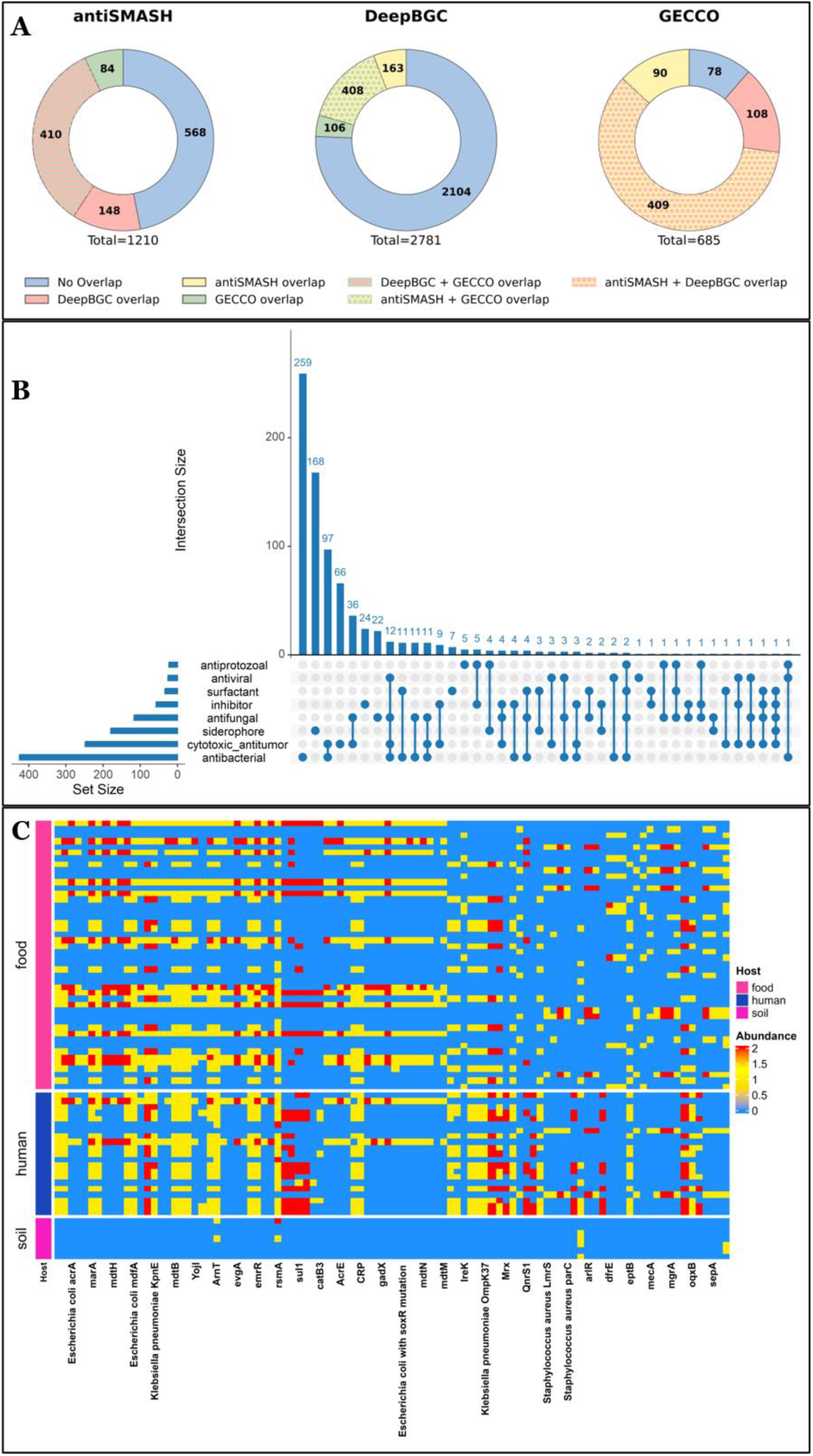
Multipanel figure NCBI dataset (A) BGC prediction overlap across tools in the NCBI dataset (B) An upset plot showing the overlap of predicted bioactivities of BGC and (C) A heatmap presenting the top 10% prevalent AMR genes distribution across habitat. Donut plots showing the overlap of BGCs predicted by antiSMASH, DeepBGC, and GECCO in the NCBI dataset. Each segment represents either unique predictions (no overlap) or shared predictions between tools (pairwise or multi-tool overlaps). Total BGC counts per tool are indicated below each plot. The vertical bars represent the shared bioactivities of BGCs annotated from the NCBI dataset.

Further, to strengthen confidence in predicted bioactivities, NPBdetect and NPF prediction overlap analysis was performed, revealing that 298 BGCs were predicted to be antibacterial, followed by 49 BGCs predicted to be cytotoxic/antitumor, and 36 BGCs predicted to be antifungal by both prediction tools. This provides greater confidence in the robustness of these bioactivity predictions (Supplementary Figure 2B).

### Characterization of predicted siderophore BGCs

Out of 180 BGCs with predicted probabilities ≥0.5 for siderophore bioactivity, 161 BGCs were found to encode at least one of the key siderophore-associated transporter domains, namely TonB, FecCD, and PBP. Analysis of the distribution of these domains, identified through independent analysis, revealed both distinct and overlapping occurrence patterns across the dataset. The most prevalent architecture comprised the simultaneous presence of all three domains, observed in 86 BGCs (53.4%), indicating that more than half of the predicted siderophore BGCs encode a relatively complete transport and uptake machinery (Supplementary Figure 2C). In addition, 30 BGCs (18.6%) contained the FecCD–PBP combination, while 4 BGCs (2.5%) encoded FecCD–TonB, and only 1 BGC (0.6%) encoded the PBP–TonB combination. Single-domain occurrences were comparatively rare for FecCD (3 BGCs; 1.9%) and PBP (3 BGCs; 1.9%), whereas TonB alone was detected in 34 BGCs (21.1%). Collectively, these findings suggest that the majority of predicted siderophore BGCs in the NCBI use case possess a more integrated transporter architecture, with fewer clusters exhibiting partial domain combinations.

Analysis of antiSMASH siderophore-associated detection rules showed that 154 clusters were assigned to known siderophore-associated rule categories, while 26 BGCs did not match any established antiSMASH siderophore rule. Among the classified clusters, NRP-metallophore-associated pathways were the most abundant, accounting for 68.8%, followed by NI-siderophore pathways at 30.5%. A very small fraction (0.6%) was associated with aminopolycarboxylic acid-like siderophore biosynthesis (Supplementary Figure 2D). Overall, these results indicate that the predicted siderophore BGCs in the NCBI dataset are dominated by NRPS-dependent metallophore-like architectures, while a subset of clusters remains unclassified by current antiSMASH siderophore rule definitions, potentially reflecting atypical or divergent siderophore-associated biosynthetic systems.

Further, the clustering of predicted siderophore BGCs with reference NRP-metallophore BGCs revealed the clustering of 27 predicted BGCs with reference metallophore BGCs, highlighting the presence of three core siderophoric moieties, such as catechol, OHAsp, and hydroxamate, and salicylate associated with these siderophoric BGCs (Supplementary Table 6).

### Profiling and surveillance of AMR gene dissemination using BGX pipeline

As mentioned previously, genomes isolated from soil, food, and human samples were analysed using CARD-RGI to detect resistance gene distribution in the IndeNCBI dataset. Likewise, 546 resistance genes were detected, and the number of hits per gene was recorded. We marked a notable concordance in the resistance profile of food and human samples (Supplementary Figure 2E). An overlap of 38.2% (n=113) resistance genes was observed. However, human samples and soil samples share the second largest overlap of 18.2% (n= 54). It was noteworthy to mention that samples collected from human sources still contain the highest percentage of unique genes (18.2%, n=54). Comparably, food had the second-highest number of unique resistance genes, 14.9% (n=44), followed by soil, 7.1% (n=21) (Supplementary Figure 2F). However, soil-to-food transmission remained quite low, with an overlap of 1.4% (n=4), and only 2% (n=6) genes were common across three sources. We have listed the top ten most prevalent AMR genes across habitats - *rsmA*, *ArnT,* Haemophilus influenzae PBP3 conferring resistance to beta-lactam antibiotics, *acrB*, *msbA*, Escherichia coli mdfA, Klebsiella pneumoniae *kpnF*, Klebsiella pneumoniae *kpnE, mdtB, and mdtC* (Figure 5C). The spread of several β-lactam antibiotics, including *TEM-1*, *CTX-M-15*, efflux pump genes including *oqxA* and *oqxB,* and other multidrug resistance genes, including *sul1*, *ompA,* across different habitats warrants further investigation on the spread of multidrug resistance genes (Figure 5C). As reported before, soil and contaminated foods are the biggest reservoirs of AMR genes and a major source of dissemination (Das et al., 2025; Islam et al., 2025; Read et al., 2024; Sassi et al., 2025). We observed that 38.2% of resistance genes were shared between food and human samples and 18.2% were shared between human and soil samples, highlighting a major passage of resistance gene transfer. Despite the considerable overlap, 18.2% of resistance genes were found to be unique to human samples in the IndeNCBI dataset. This observation underscores the need for further investigation into potential contributions from other habitats, along with the impact of over-the-counter drug access and the misuse of antibiotics. Consistent with the index case, IndeNCBI dataset analysis revealed that *rsmA* is the most prevalent resistance gene. The gene rsmA encodes a post-transcriptional regulatory protein that modulates multiple key regulators of virulence-associated functions, including biofilm formation (Mulcahy et al., 2008). Likewise, in the ODOG dataset, arnT emerged as the second most prevalent AMR gene in the IndeNCBI dataset. This is particularly concerning, as it is known to confer resistance to polymyxin antibiotics (Sasal et al., 2025). Prevalence of *sul1* and *qacEdelta1* resistance genes was previously reported as prevalent ARG in a surveillance study conducted in Japan (Sekizuka et al., 2022). This emphasizes the BGX pipeline’s role in genomics-based surveillance for tracking the spread of resistance genes from the environment to humans.

## Conclusion

Across both use cases, the BGX pipeline consistently demonstrates its ability to capture diverse BGCs, predict the bioactivity of BGCs, and determine AMR gene dissemination with high confidence as compared to other workflows (Supplementary Table 7). The dataset from the ODOG initiative shows a highly heterogeneous habitat distribution with strong environmental representation, whereas the IndeNCBI dataset is comparatively more balanced but skewed toward food and host-associated sources. Multi-tool integration for BGC prediction highlights clear methodological complementarity: DeepBGC provides high sensitivity, while antiSMASH and GECCO contribute specificity, and their overlaps define a robust set of high-confidence BGCs. The ODOG initiative dataset shows a greater proportion of unique predictions, indicating higher novelty, while the IndeNCBI dataset exhibits greater overlap, reflecting conserved and well-characterized biosynthetic regions. ODOG contains a substantial fraction of novel and diverse BGCs, particularly enriched in terpene and RiPP classes, indicating significant unexplored biosynthetic potential. In contrast, the IndeNCBI dataset is dominated by KNOWN_LIKE clusters, with novelty largely restricted to terpene biosynthesis, suggesting a more conserved metabolic landscape. The integrated bioactivity prediction across ODOG and IndeNCBI datasets reveals a rich reservoir of bioactive BGCs, with antibacterial activity being the most dominant. Overlap-based analysis enabled the identification of high-confidence antibacterial, cytotoxic, and antifungal clusters, enhancing prediction reliability. Comparative trends indicate a predominance of NRPS-independent siderophore pathways in ODOG, while IndeNCBI is enriched in NRPS-dependent metallophore systems. Transporter domain analysis shows that many siderophore BGCs encode complete uptake machinery, especially in the IndeNCBI dataset, suggesting functional robustness. The presence of conserved siderophoric moieties further validates the biological relevance of predictions. Additionally, unclassified clusters highlight the potential for novel biosynthetic pathways. Overall, this approach provides a robust framework for prioritizing high-confidence BGCs for natural product discovery. The antimicrobial resistance gene overlaps between the environment and human samples in both the ODOG and the IndeNCBI datasets underscore the environment as a major driver of ARGs transmission. The prevalence of *rsmA* and *arnT* makes them key resistance determinants across datasets. Notably, a significant fraction of human-specific ARGs warrants further investigation of underexplored sources. This underscores the applicability of this pipeline with openly available data to track the emergence and transmission of multidrug resistance genotypes. In the ODOG dataset we found 10 C-type lectins (Supplementary Table 8) and in the IndeNCBI dataset we found 6 C-type lectins (Supplementary Table 9). The C-type lectin belonged to diverse habitats like marine, gut of fresh water puffer fish, urine, renal tubules and genital tract, human upper respiratory tract, human, wastewater, skin and rice. The C-type lectins recognise specific glycans and participate in host-pathogen interactions and cell adhesion (Brown et al., 2018). Finding C-type lectins in the bacterial genomes suggests calcium-based carbohydrate-binding capabilities.

As a standalone feature, BGX enables the export and visualization of all generated data in a separate HTML file for easy interpretation. A variety of visualization techniques were employed to enhance data representation, including bar charts, donut plots, UpSet plots, and heatmaps. Additionally, the original CSV files were provided to enable further analysis by the user. Availability of this pipeline in a Docker container meets the challenge of installing different tools and recurring changes of environment. Once installed, the utility scripts inside the Docker container make it adaptable for specific analyses, such as genomic surveillance. Future developments will expand the pipeline coverage from genome to metagenome and metagenome-assembled genomes. New tools will be incorporated to elucidate novel features, machine learning-based methods to draw new inferences, and AI-powered search functionalities to further enrich the breadth and diversity of BGX.

## Data and Code availability

All analyzed data, processed outputs with high resolution figures, and supplementary files generated during this study are available in the link given below: https://drive.google.com/drive/folders/1zhma822KVrWps7pMOz5-rVVpPBQOod4q?usp=sharing

We welcome collaborations from researchers and organizations interested in utilizing, evaluating, improving, or further developing the BGX pipeline and its associated resources. For further details and to collaborate on our pipeline, please visit the official webpage of BGX at: https://bgx.nabi.res.in

## Author’s contribution

AC, LS, BK, and SP contributed equally to this work. SM conceptualized and designed the study. BK curated the data. AC, LS, BK, and SP did the formal bioinformatic analysis. BK and LS developed the BGX pipeline, and BK and AC tested and validated the pipeline. LS developed the interface, and BK created the Docker container for pipeline deployment. AC, LS, BK, and SP wrote the manuscript. SM supervised and reviewed the manuscript. All the authors reviewed, edited, and approved the final version.

## Funding

This work was supported by BRIC - National Agri-Food and Biomanufacturing Institute Core Funding (2026) to Shrikant Mantri.

## Competing Interest

The authors declare no conflict of interest.

## Acknowledgements

We would like to thank BRIC-National Agri-Food and Biomanufacturing Institute (NABI) for providing the necessary infrastructure and supporting high-performance computing facilities. We acknowledge the National Supercomputing Mission (NSM) for providing computing resources of PARAM Smriti at BRIC-NABI, Mohali, which is implemented by C-DAC and supported by the Ministry of Electronics and Information Technology (MeitY), Department of Biotechnology (DBT), and Department of Science and Technology (DST), Government of India. We also acknowledge DeLCON (DBTelectronic library consortium), Gurugram, India, for the online journal access. SP is thankful to Regional Center of Biotechnology (RCB), Department of Biotechnology, Government of India (GOI) for undertaking PhD registration. SM and BK would like to acknowledge NABI Core funding for the execution of this project. SM, AC, and LS acknowledge the financial support provided by the Department of Biotechnology (DBT), Ministry of Science and Technology, Government of India, through project grant BT/PR54729/BSA/33/250/2024 and the BRIC-NABI Biomanufacturing BioFoundry project “Agri-Food Bio-Mug” (BT/TEMP/24019/BMH-01/24). The authors thank Manvinder Jaswal for his contribution to the conceptualization of the introduction.

We acknowledge the Biotechnology Research and Innovation Council (BRIC) One Day One Genome (ODOG) initiative, coordinated by DBT and BRIC–National Institute of Biomedical Genomics (BRIC-NIBMG), for providing a valuable use case that facilitated the development, testing, and demonstration of the BGX pipeline.

## Notes

https://drive.google.com/drive/folders/1zhma822KVrWps7pMOz5-rVVpPBQOod4q?usp=sharing

### Competing Interest Statement

The authors have declared no competing interest.

