## Supplemental figure for "BGX: A Comprehensive Pipeline for Genomic Insight into Bioactivity Prediction, Genomic Surveillance, and Novel Biosynthetic Gene Cluster Assessment"

### List of Supplementary Figures


| Name | Description | Page No. |
|---|---|---|
| Supplementary Figure 1A | BUSCO-based genome assessment and category distribution of ODOG initiative genomes | 2 |
| Supplementary Figure 1B | Shared bioactivities and high-confidence BGCs in the ODOG dataset | 3 |
| Supplementary Figure 1C | Venn diagram illustrating the co-occurrence of key siderophore-associated transporters in the ODOG dataset | 4 |
| Supplementary Figure 1D | Distribution of antiSMASH siderophore-associated detection rules among predicted siderophore BGCs in the ODOG dataset | 5 |
| Supplementary Figure 1E | AMR gene distribution across wastewater, human samples, and soil from ODOG data | 6 |
| Supplementary Figure 1F | Distribution of resistance genes across multiple habitats within the ODOG dataset. | 7 |
| Supplementary Figure 2A | BUSCO-based genome assessment and category distribution of the IndeNCBI dataset | 8 |
| Supplementary Figure 2B | Shared bioactivities and high-confidence BGCs in the IndeNCBI dataset | 9 |
| Supplementary Figure 2C | Venn diagram illustrating the co-occurrence of key siderophore-associated transporters in the IndeNCBI dataset | 10 |
| Supplementary Figure 2D | Distribution of antiSMASH siderophore-associated detection rules among predicted siderophore BGCs in the IndeNCBI dataset | 11 |
| Supplementary Figure 2E | AMR gene distribution across food, human samples, and soil from IndeNCBI data | 12 |
| Supplementary Figure 2F | Distribution of resistance genes across multiple habitats within the IndeNCBI dataset | 13 |

**Supplementary Figure 1A -** BUSCO-based genome assessment and category distribution of ODOG initiative genomes

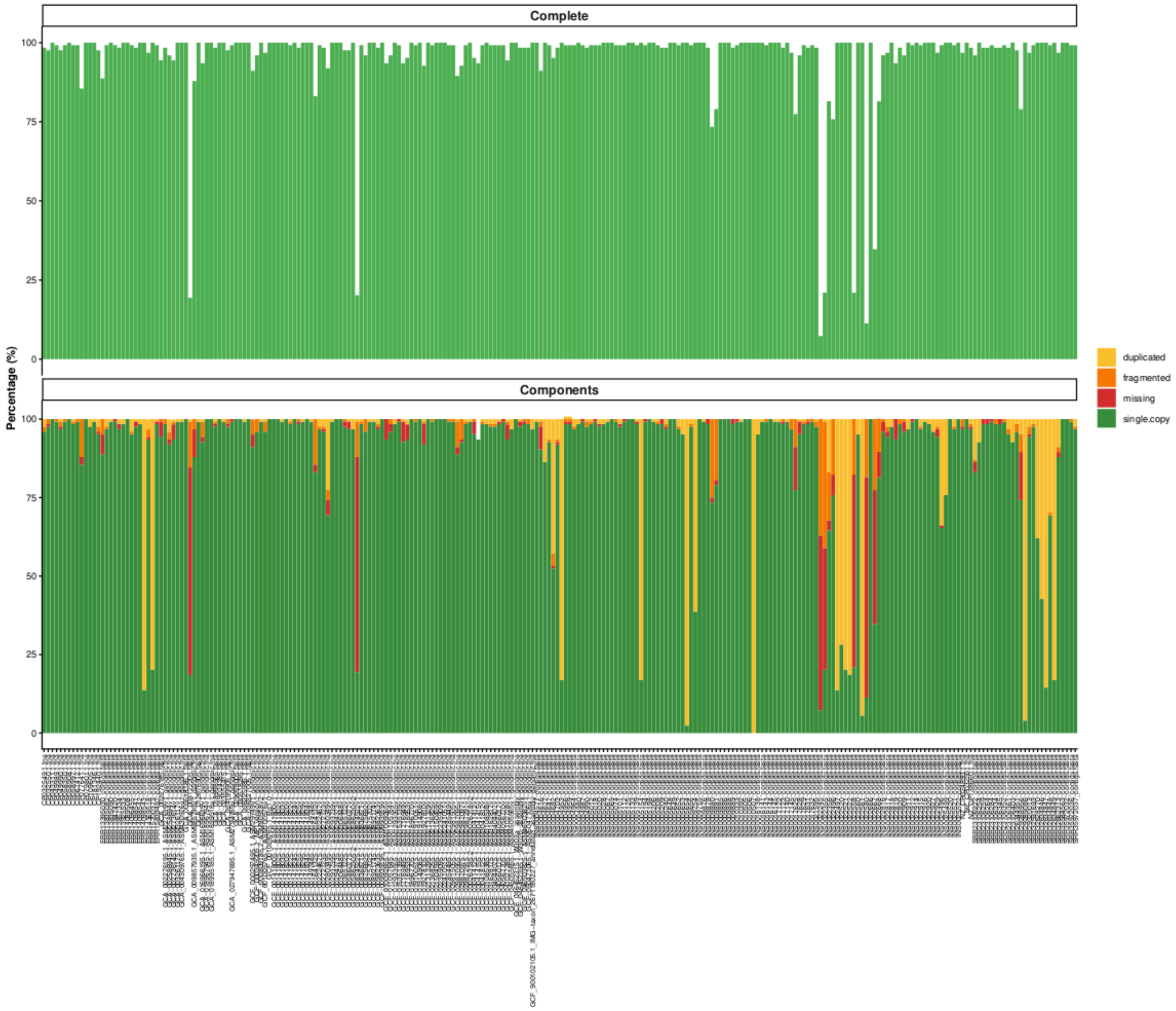


Figure 1A. BUSCO-based genome assessment and category distribution of ODOG initiative genomes. . The distribution of complete (single-copy and duplicated), fragmented, and missing categories across ODOG genomes provides an overview of genome quality and highlights the proportion of high-quality genomes with predominantly complete BUSCOs.

**Supplementary Figure 1B -** Shared bioactivities and high-confidence BGCs in the ODOG dataset

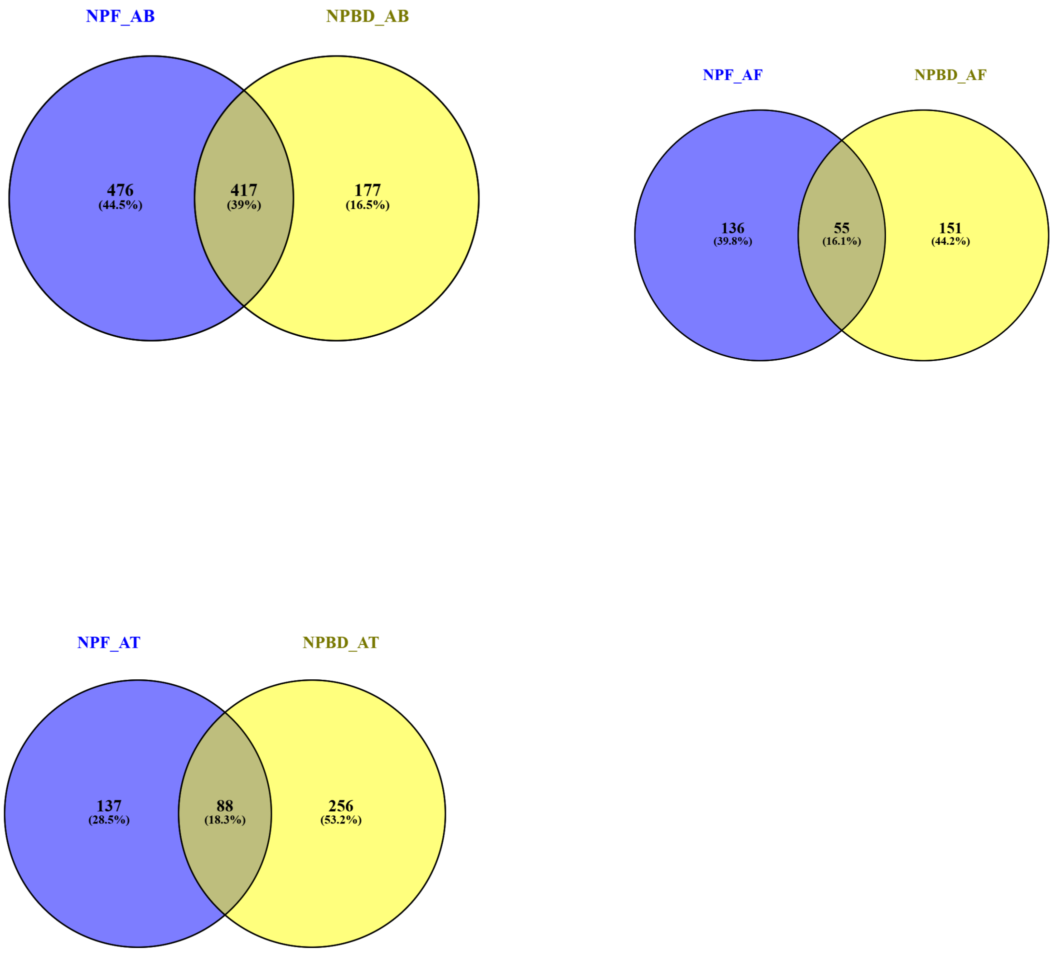


Figure 1B. Venn diagram showing the overlap among predicted antibacterial (AB), antifungal (AF), and cytotoxic/antitumor (AT) BGCs identified from ODOG use case. The diagram highlights BGCs with shared predicted bioactivities and identifies high-confidence clusters supported by overlapping bioactivity predictions.

**Supplementary Figure 1C -** Venn diagram illustrating the co-occurrence of key siderophore-associated transporters in the ODOG dataset

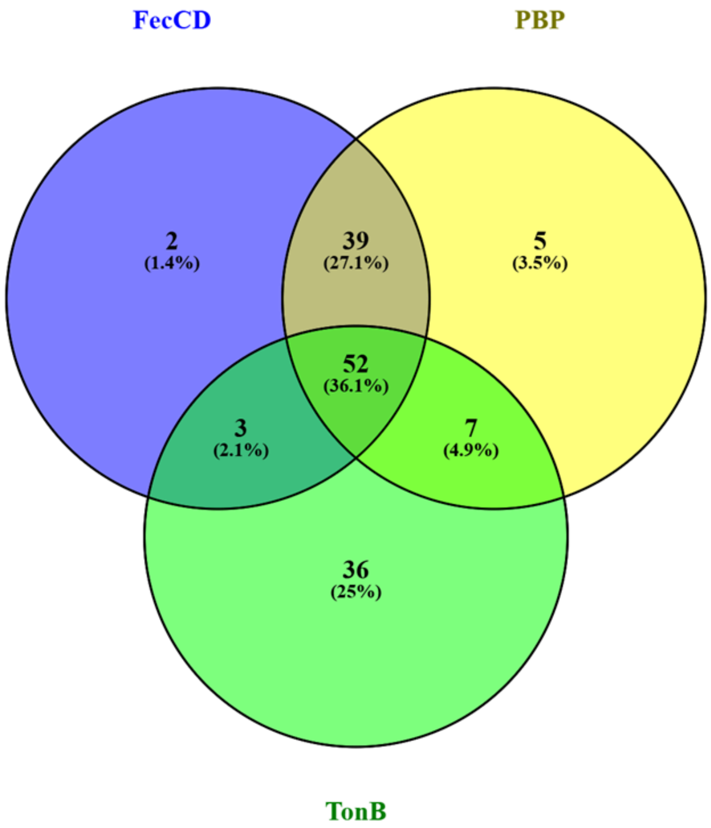


Figure 1C: Venn diagram representing the occurrence and co-occurrence of key siderophore-associated transporter domains (FecCD, PBP, and TonB) in 144 predicted siderophore BGCs identified in the ODOG use case.

**Supplementary Figure 1D -** Distribution of antiSMASH siderophore-associated detection rules among predicted siderophore BGCs in the ODOG dataset

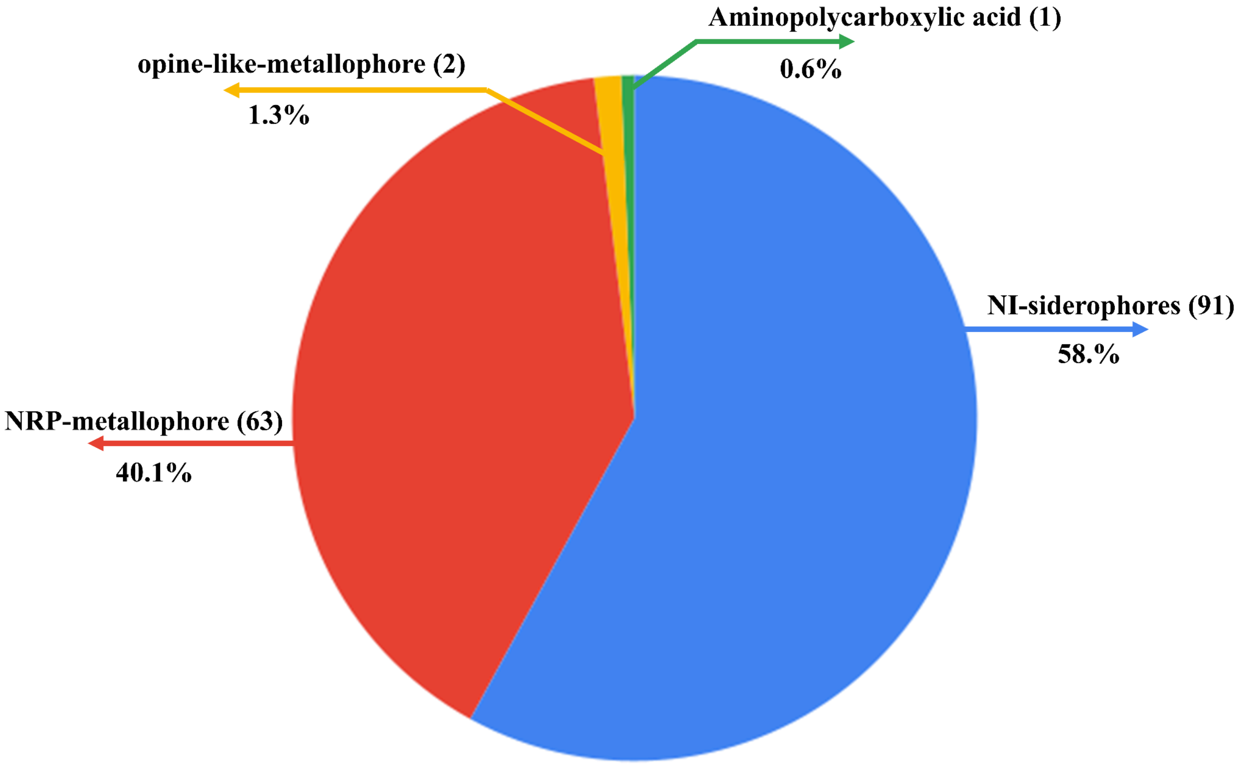


Figure 1D: Pie chart illustrating the distribution of antiSMASH siderophore-associated detection rules among the predicted siderophore BGCs.

**Supplementary Figure 1E -** AMR gene distribution across wastewater, human samples, and soil from the ODOG data

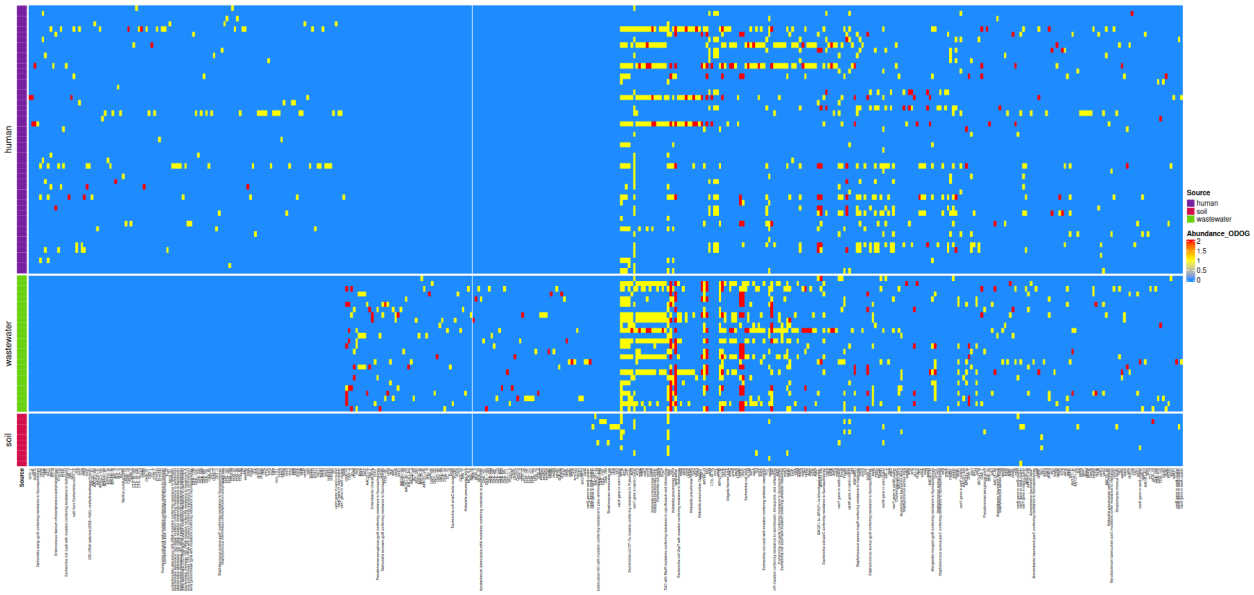


Figure 1E. AMR gene distribution across soil, wastewater, and human samples from the ODOG dataset. The figure illustrates the presence and relative abundance of AMR genes across different environments and habitats, providing insights into the dissemination and ecological distribution of AMR genes.

**Supplementary Figure 1F -** Distribution of resistance genes across multiple habitats within the ODOG dataset.

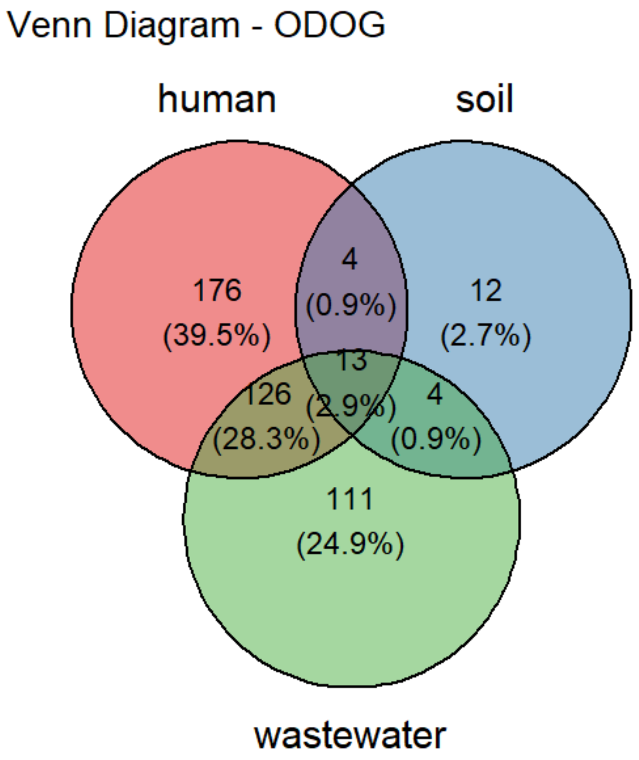


Figure 1F. Resistance genes are shared across different habitats of ODOG initiative dataset.

**Supplementary figure 2A -** BUSCO-based genome assessment and category distribution of the IndeNCBI dataset

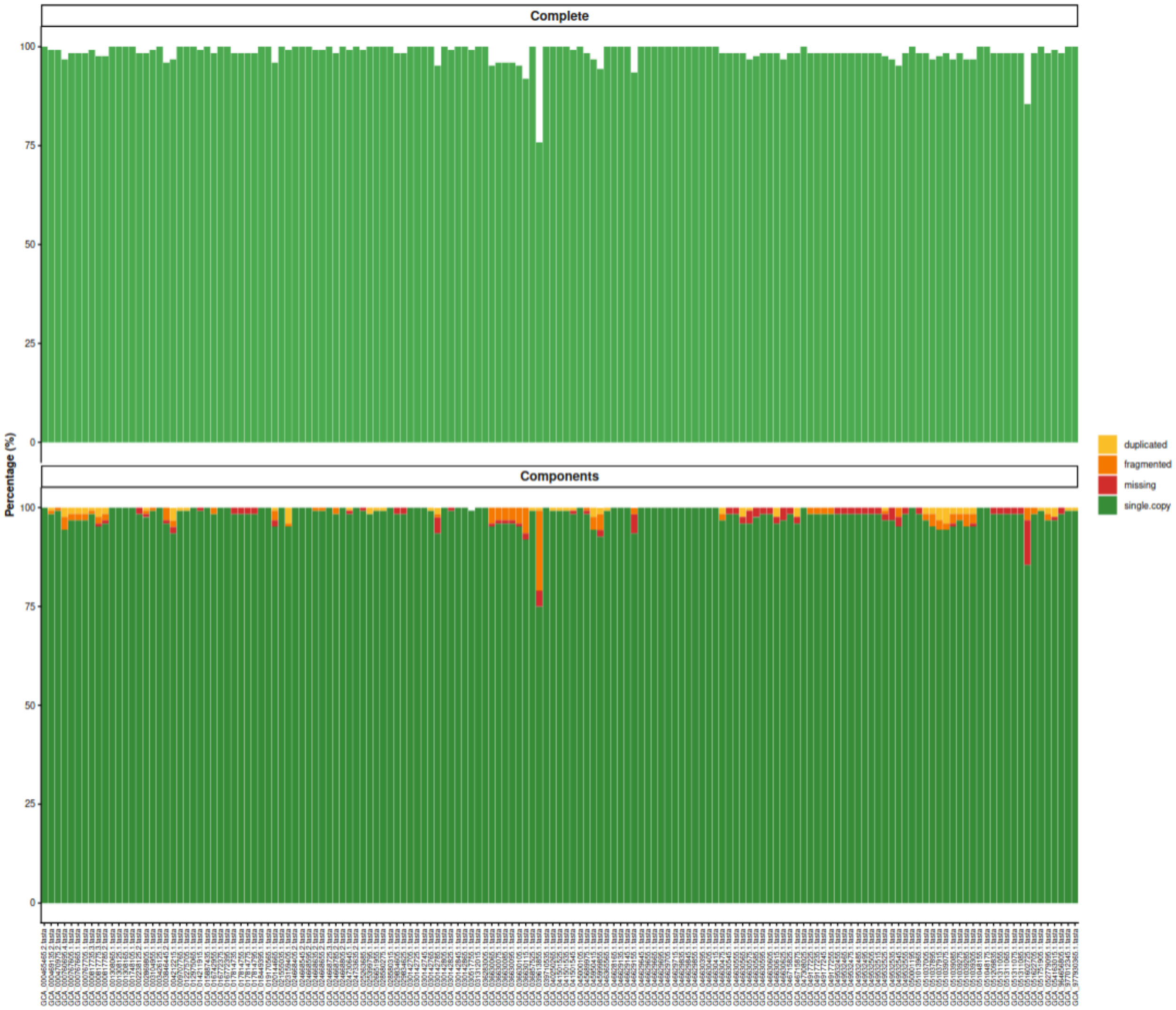


Figure 2A. BUSCO-based genome assessment and category distribution of the IndeNCBI genomes. The distribution of complete (single-copy and duplicated), fragmented, and missing categories across the dataset provides an overview of genome quality and highlights the proportion of high-quality genomes with predominantly complete BUSCOs.

**Supplementary Figure 2B -** Shared bioactivities and high-confidence BGCs in the IndeNCBI dataset

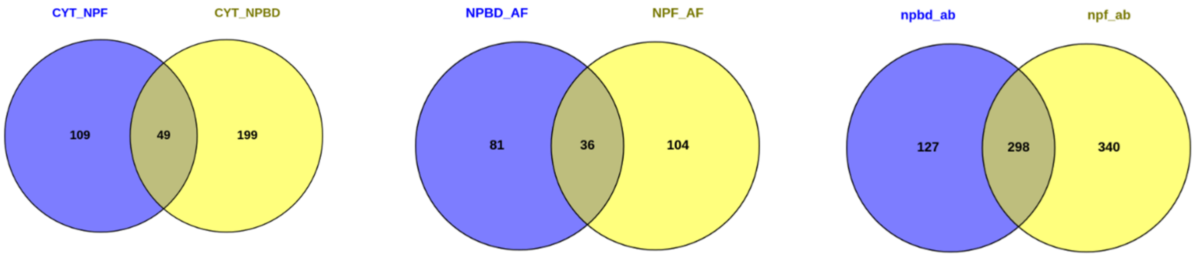


Figure 2B. Venn diagram showing the overlap among predicted antibacterial (ab), antifungal (AF), and cytotoxic/antitumor(CYT) BGCs identified from the IndeNCBI use case. The diagram highlights BGCs with shared predicted bioactivities and identifies high-confidence clusters supported by overlapping bioactivity predictions.

**Supplementary Figure 2C -** Venn diagram illustrating the co-occurrence of key siderophore-associated transporters in the IndeNCBI dataset

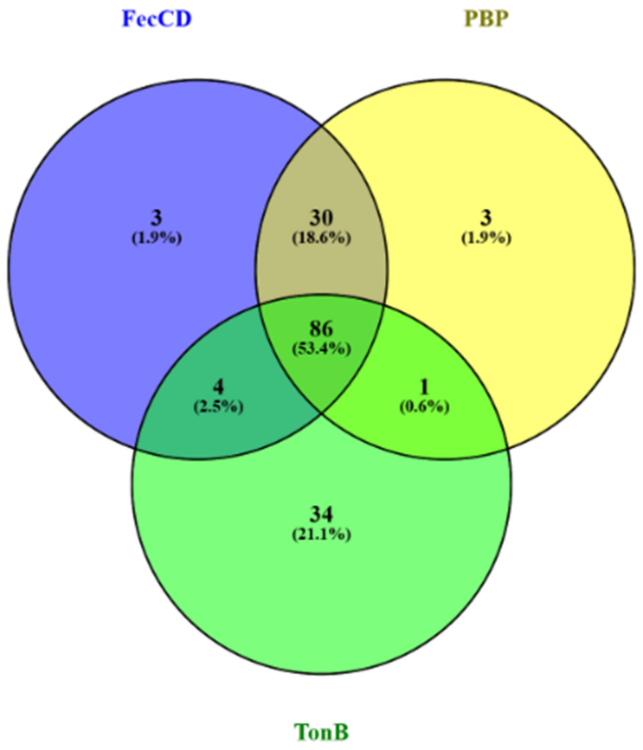


Figure 2C: Venn diagram representing the occurrence and co-occurrence of key siderophore-associated transporter domains (FecCD, PBP, and TonB) among 161 predicted siderophore biosynthetic gene clusters (BGCs) identified in the NCBI use case

**Supplementary Figure 2D -** Distribution of antiSMASH siderophore-associated detection rules among predicted siderophore BGCs in the IndeNCBI dataset

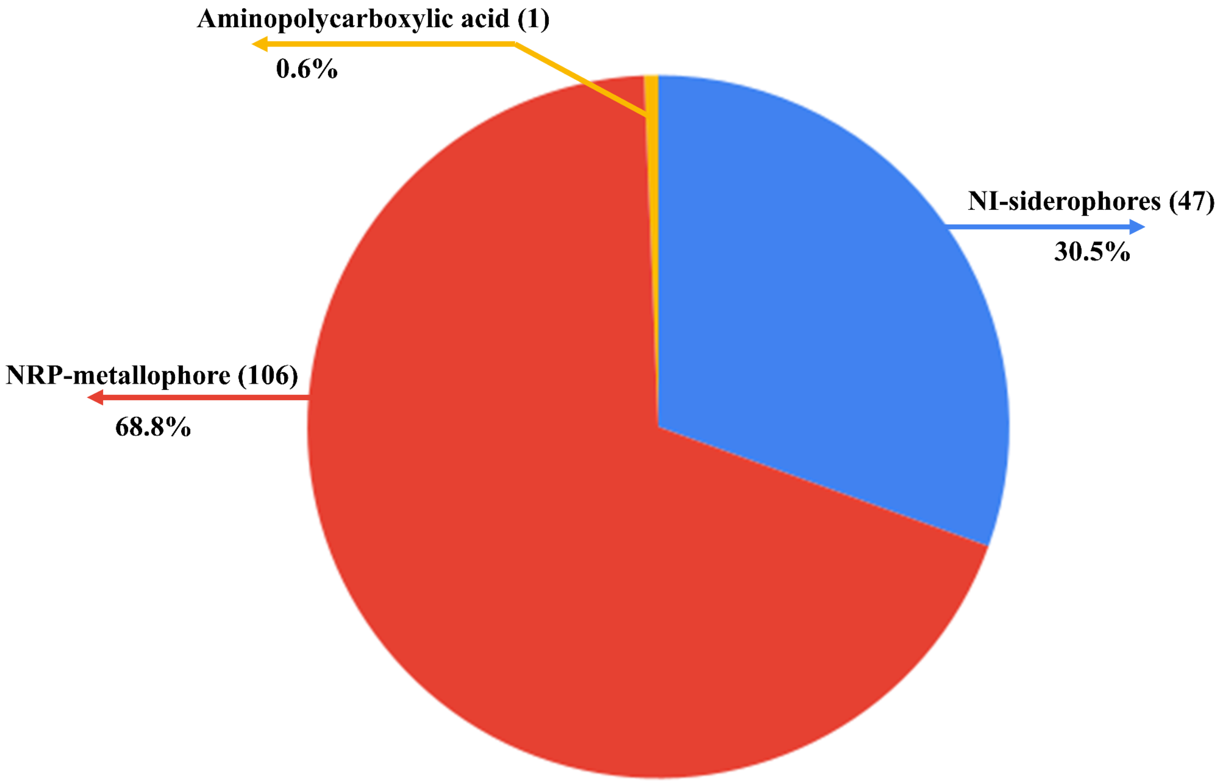


Figure 2D. Pie chart showing the distribution of antiSMASH siderophore-associated detection rules among the predicted siderophore BGCs in the NCBI use case, highlighting the predominance of NRP-metallophore and NI-siderophore rule categories.

**Supplementary Figure 2E** - AMR gene distribution across food, human samples, and soil from IndeNCBI data

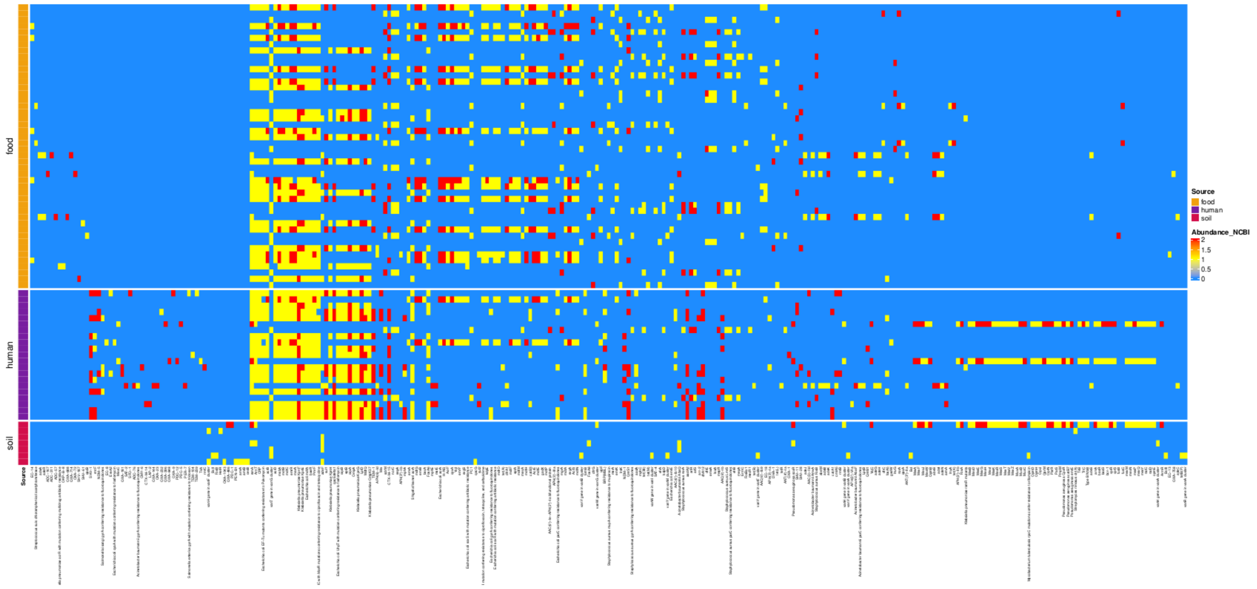


Figure 2E. AMR gene distribution across soil, human, and food samples from the IndeNCBI dataset. The figure illustrates the presence and relative abundance of AMR genes across different environments and habitats, providing insights into the dissemination of AMR genes.

**Supplementary Figure 2F -** Distribution of resistance genes across multiple habitats within the IndeNCBI dataset.

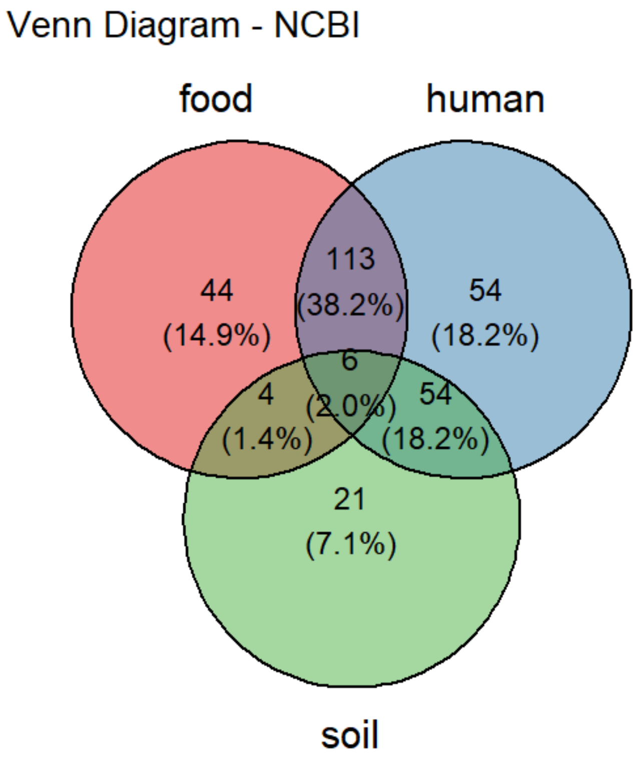


Figure 2F. Resistance genes are shared across different habitats of the IndeNCBI dataset.
