## Supplemental table for "BGX: A Comprehensive Pipeline for Genomic Insight into Bioactivity Prediction, Genomic Surveillance, and Novel Biosynthetic Gene Cluster Assessment": Supplementary tables index.pdf

### List of Supplementary Tables

| Name | Description |
|---|---|
| Supplementary Table 1A | Metadata ODOG use case |
| Supplementary Table 1B | Summary of BGC Predictions from Multiple Tools-ODOG |
| Supplementary Table 1C | BiG-SCAPE clustering summary of antiSMASH 8-annotated BGCs from the ODOG use case. |
| Supplementary Table 1D | Annotation of antiSMASH predicted BGCs-ODOG dataset |
| Supplementary Table 2 | BA prediction of the ODOG use case |
| Supplementary Table 3 | Siderophoric moieties associated with predicted siderophore BGCs in the ODOG dataset, ref gcf information |
| Supplementary Table 4A | Metadata IndeNCBI use case |
| Supplementary Table 4B | Summary of BGC Predictions from Multiple Tools-IndeNCBI dataset |
| Supplementary Table 4C | BiG-SCAPE clustering summary of antiSMASH 8-annotated BGCs from IndeNCBI use case |
| Supplementary Table 4D | Annotation of antiSMASH predicted BGCs-IndeNCBI dataset |
| Supplementary Table 5 | BA prediction of the IndeNCBI use case |
| Supplementary Table 6 | Siderophoric moieties associated with predicted siderophore BGCs in the IndeNCBI dataset |
| Supplementary Table 7 | Feature comparison table |
| Supplementary Table 8 | C-type lectins found in the ODOG dataset |
| Supplementary Table 9 | C-type lectins found in the IndeNCBI dataset |
